# Accounting for observation biases associated with counts of young when estimating fecundity: case study on the arboreal-nesting red kite (*Milvus milvus*)

**DOI:** 10.1101/2023.12.01.569571

**Authors:** Sollmann Rahel, Adenot Nathalie, Spakovszky Péter, Windt Jendrik, Mattsson Brady J.

**Affiliations:** Department of Ecological Dynamics, Leibniz Institute for Zoo and Wildlife Research – Berlin, Germany; Institute of Wildlife Biology and Game Management, Department of Integrative Biology and Biodiversity Research, University of Natural Resources and Life Sciences – Vienna, Austria; TB Raab GmbH – Deutsch-Wagram, Austria

**Keywords:** false positives, false negatives, multinomial model, classification, calibration, population dynamics

## Abstract

Counting the number of young in a brood from a distance is common practice, for example in tree-nesting birds. These counts can, however, suffer from over and undercounting, which can lead to biased estimates of fecundity (average number of nestlings per brood). Statistical model development to account for observation bias has focused on false negatives (undercounts), yet it has been shown that these models are sensitive to the presence of false positives (overcounts) when they are not accounted for. Here, we develop a model that estimates fecundity while accounting for both false positives and false negatives in brood counts. Its parameters can be estimated using a calibration approach that combines uncertain counts with certain ones, which can be obtained by accessing the brood, for example during ringing. The model uses multinomial distributions to estimate the probabilities of observing *y* young conditional on the true state of a brood *z* (i.e., true number of young) from paired uncertain and certain counts. These classification probabilities are then used to estimate the true state of broods for which only uncertain counts are available. We use a simulation study to investigate bias and precision of the model and parameterize the simulation with empirical data from 26 red kite nests visited with ground and nest-based counts during 2021 and 2022 in central Europe. In these data, bias in counts was at most 1 in either direction, more common in larger broods, and undercounting was more common than overcounting. This led to an overall 5% negative bias in fecundity in uncertain counts. The model produced essentially unbiased estimates (relative bias < 2%) of fecundity across a range of sample sizes. This held true whether or not fecundity was the same for nests with paired counts and those with uncertain-only counts. But the model could not estimate parameters when true states were missing from the paired data, which happened frequently in small sample sizes (n = 10 or 25). Further, we projected populations 50 years into the future using fecundity estimates corrected for observation biases from the multinomial model, and based on “raw” uncertain observations. We found that ignoring observation bias led to strong negative bias in projected population size for growing populations, but only minor negative bias in declining populations. Accounting for apparently minor biases associated with ground counts is important for ensuring accurate estimates of abundance and population dynamics especially for increasing populations. This could be particularly important for informing conservation decisions in projects aimed at recovering depleted populations.

## Introduction

Understanding spatiotemporal mechanisms for changes in animal abundance rests on information about survival, movement, and fecundity (Sample et al. 2018, Sen and Akçakaya 2022). Challenges when estimating these parameters due to observation bias are notorious (Kéry and Royle 2015), and correcting for these observation biases requires the use of statistical methods such as capture-recapture models when estimating survival, recruitment and/or abundance (Williams et al. 2002). Capture-recapture models also serve as a core building block of integrated population models (Zipkin and Saunders 2018, Zhao 2020). Several quantities have been proposed to measure fecundity (Etterson et al. 2011). Existing methods for estimating fecundity, however, typically do not address inaccurate counts of offspring (but see Sen and Akçakaya 2022). Failing to account for such bias may undermine inferences about population dynamics and their drivers at this critical part of the annual life cycle, and in turn compromise efforts to conserve species.

A particular challenge when estimating fecundity is ascertaining the number of young produced per litter or per nest (henceforth, brood). The number of young can be determined accurately by accessing the brood close to fledging in altricial species, as is often done in bird ringing programs (e.g., Katzenberger et al. 2019). Depending on nest location, doing so may require specialized training and equipment (e.g., climbing or using a lifting platform for tree-nesting birds), is time-intensive (Fuller and Mosher 1981) and can stress and disturb the adults (Andersen 1990), which could reduce productivity (Ibáñez-Alamo et al. 2012). In some cases, accessing a nest may not be possible at all if the structure supporting the nest is too fragile (e.g. a thin branch that cannot support human weight). To minimize disturbance, costs, risk to the observer, and simplify field logistics, counts of young are therefore often obtained at a distance from the ground, for example in tree-nesting raptors (Brown et al. 2013). Nest monitoring can also be done from the air using helicopters (White and Sherrod 1973) or more recently drones (Gallego and Sarasola 2021). In this context, counts of young may not reflect true brood size, and obtaining an accurate count likely depends on several factors, including observation distance, time spent observing, level of camouflage, activity level of the young as well as their position in the nest (i.e., central or close to the edge).

Determining brood size, when done via observation from a distance, potentially suffers from two types of observation error, false negatives (when young are missed; e.g., Goszczyński 1997) and false positives (when young are double-counted, as occurred in our surveys; see below). False negatives have long been acknowledged as pervasive in wildlife surveys, and there is a wealth of statistical models dealing with false negatives: capture-recapture models (Otis et al. 1978), N-mixture (Royle 2004), or distance sampling models (Buckland et al. 2015) estimate abundance when the probability of detecting any individual is < 1; and occupancy models (MacKenzie et al. 2002, Tyre et al. 2003) estimate occurrence when species detection is < 1. Even though several studies have shown that false positives are also common in wildlife surveys (Clement et al. 2014, Brack et al. 2018, McKibben et al. 2023) and that models accounting only for false negatives are very sensitive to the presence of false positives (Miller et al. 2011, Lahoz-Monfort et al. 2016, Link et al. 2018, Nakashima 2020), the development of models accounting for false positives has lagged behind (Dénes et al. 2015). Existing efforts to formally account for false positives have largely focused on occupancy models (Royle and Link 2006, Miller et al. 2011, Chambert et al. 2015), which require some form of additional information to be uniquely identifiable, including constraints on parameters (Royle and Link 2006) or detections with and without false positives (Miller et al. 2011, Chambert et al. 2015). In the context of abundance estimation, the Poisson-Poisson version of the N-mixture model (Kéry and Royle 2015) allows for counting an individual more than once but requires repeated visits. In contrast, here, we develop and test a method that accounts for both over and undercounting based on a single visit to each nest. We are unaware of any studies that have accounted for false positive bias when estimating brood size.

To do so, we treat brood size as a state that is imperfectly observed when the young are counted from a distance. We then expand the approach of Royle and Link (2006) to formulate observation state probabilities (here, observed number of young) as conditional on true states (here, true number of young) to >2 states. Without additional information, parameters in this model are not uniquely identifiable, and we adopt a modified version of the calibration approach by Chambert et al. (2015), in which for some sites the true state is confirmed (here, nests that are climbed and young counted without error) to inform the imperfect observation process. We evaluate the performance of this model using a case study on the red kite (*Milvus milvus*) in central Europe. Finally, we investigate the implications of ignoring observation error in brood size counts for accuracy in predicted population trajectories.

## Methods

### Empirical data collection

We collected data on red kite brood size during June of 2021 and 2022 during the fledging period. From the middle of June onwards and approximately three weeks after hatching, young reach a development phase that is sufficient for tagging and ringing (Cramp and Simmons 1980). A total of 26 nests were identified from the ground at the beginning of the breeding season by members and partners of the LIFE EUROKITE (LIFE18 NAT/AT/000048) project, and checked again at the end of June to confirm occupancy. In 2021, we sampled 8 nests in Germany. In 2022, 14 of the 18 sampled nests were located in stands of deciduous forest within Lower Austria, along the March River that forms the border with Slovakia. Remaining nests that year (i.e., 3 in Upper Austria and 1 in Slovakia) were located in mixed deciduous-coniferous forests.

Each nest was visited once by two observers. The ground observer aimed to find the best vantage point between 50 m and 100 m from the nest, and used a spotting scope or binoculars to observe nestlings. Counting from the ground proceeded before and during climbing of the tree by the second observer, a person certified for tree-climbing along with tagging and ringing of the young red kites. The ground observation started as soon as the best vantage point to observe the nest was found and stopped when the climber reached the nest. The climber did not reveal the number of young in the nest until after the ground-count was conducted.

### Model development

Royle and Link (2006) developed an occupancy model allowing for false positive and false negative detections by formulating probabilities of observed states *y* (0 or 1) conditional on true states *z* (0 or 1). For example, when a site is not occupied (*z* = 0), we can either obtain a true negative observation (*y* = 0) with probability *p*, or we can obtain a false positive observation (*y* = 1) with probability 1 − *p*. They conceptually expanded this model to more than 2 states; for each true state, the probabilities of all possible observed states sum to 1, and observations *y* for a given state *z* are multinomial random variables.

Here, we adapt this approach and consider brood size as the true state with *Z* possible categories corresponding to possible numbers of nestlings. If data collection includes nests that could be empty, *z* can take values from 0 (i.e., empty nest) to the biological upper limit of how many offspring the focal species can raise. If only nests confirmed to be occupied are included, *z* = 0 is not a possible state. Regardless:

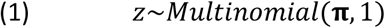

Here, **π** is the vector of probabilities that a nest falls into a given state category (henceforth, nest probabilities). The expected number of nestlings in any given nest is 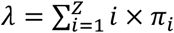. We refer to *λ* as fecundity.

Observed states, *y*, can fall into one of *K* categories (where *K* may be equal to *Z*), and the probability of observing a particular category *k, P*(*y* = *k*), depends on the true state *z*.

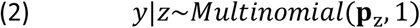

Thus, **p** is a *Z×K* classification matrix, and we refer to values in **p** as classification probabilities.

Royle and Link (2006) showed that this model is not uniquely identifiable for the 2-state and *K*-state case; further development by Miller et al. (2011) and Chambert et al. (2015) allowed for the incorporation of additional information to make parameters identifiable. Here, we adopt a modified version of the ‘calibration’ approach by Chambert et al. (2015), in which at some sites, the true state is confirmed so that direct information on the classification process is available. Specifically, we propose to combine imperfect or undertain counts *y* (e.g., ground counts of nestlings), which provide imperfect representations of the true state, with confirmed or certain counts (e.g., climb counts for raptors) which provide *z* without error. Collecting both types of data, *y* and *z*, at a set of nests allows estimation of the parameters in the classification matrix **p**. We can then use estimates of **p** to obtain estimates of the expected number of nestlings in a data set consisting only of imperfect counts, *y*.*new*. This differs from the calibration approach of Chambert et al. (2015) in that certain observations of *z* also inform the state process (ie, estimates of **π** and *λ*). We consider two possible sampling scenarios:

1. The paired counts (i.e., imperfect ground counts, *y*, with perfect climb counts, *z*; sample size *n*) and the uncertain-only counts, *y*.*new* (sample size *n*.*new*) come from the same population, i.e., they share **π** and *λ*. In this situation, one could use only measures of *z* (i.e., only the climb counts from the paired data set) to determine **π** and *λ*. But adding *y*.*new* increases sample size and thus may yield more precise estimates of these parameters. Note that in the set of formulas below, *z*.*new* is not observed, whereas for the paired data, *z* is observed.

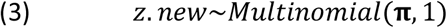

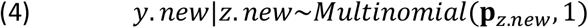
2. The paired *y* and *z* versus *y*.*new* come from different populations that may differ in fecundity (i.e., in one population, birds may produce more/fewer nestlings, on average). In this case, we can still use *y* and *z* to estimate **p** – which we assume is shared between both populations – but have to estimate a separate **π.new**.

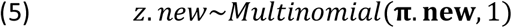

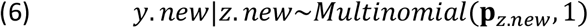

This is the more interesting scenario, as in contrast to **π, π.new** cannot be estimated from the paired data alone. Accuracy of **π.new** will depend on how accurately **p** is estimated. Depending on *Z* and *K*, the number of parameters in **p** can quickly become unwieldy (Royle and Link 2006) and require unrealistically large sample sizes to be estimated well. We address this issue in our simulation by constraining the mis-identification process such that *y* can differ from *z* by no more than 1. In the Discussion, we further present alternative model formulations that simplify the detection/classification process and reduce the number of parameters, along with some preliminary exploration of model behavior. A full implementation and presentation of these alternative formulations is beyond the scope of the current study.

### Simulation – parameter estimation

We used a simulation approach to determine how well parameters of the above described model can be estimated. We refer to data sets in which measures of both *z* and *y* from the same nests are available as ‘paired data sets’; and data sets with paired data plus additional uncertain counts *y*.*new* as ‘combined data sets’.

We parameterized our simulation based on the empirical data on the 26 paired (uncertain) ground and (certain) climb counts described above and a literature review of reproductive values in red kites. Because in our empirical data only nests with at least 1 observed nestling were included and all nests were confirmed to be occupied (i.e., no false positive observations in empty nests), we allowed *z* to take values between 1 and 4 (i.e., we did not include *z* = 0 as a possible state). Reported values of clutch size based on a global literature review ranged from 1.9 to 3.2, and number fledged per successful pair ranged from 1.4 to 2.3 (Sergio et al. 2019). As nestlings have not yet fledged when they are counted, we used reported clutch size as a guideline to set three levels of reproduction. Our baseline **π** = {0.1, 0.45, 0.4, 0.05}, resulting in fecundity *λ* = 2.4. We also created two alternative vectors of **π**, one with a higher fecundity of *λ*_*high*_ = 2.8 and a high rate of 3 nestlings: **π**_*high*_ = {0.05, 0.2, 0.65, 0.1}; one with a lower fecundity of *λ*_*low*_ = 2 and a more even distribution of nestling probabilities: **π**_*low*_ = {0.3, 0.45, 0.2, 0.05}.

Our parameterization of **p** was strongly informed by the empirical data, as to our knowledge no other published data sets for paired nestling counts are available. We assumed *K*=*Z* (implying that no observer would report more than 4 nestlings, given the knowledge of the species’ breeding habits, and because the dataset did not contain nests in which observes recorded 0 nestlings) and constructed **p** using the following rules: (1) the probability of observing the correct number of nestlings is higher than the probability of observing the wrong number of nestlings but decreases as true number of nestlings increases; (2) when observing the wrong number of nestlings, missing a nestling (false negative) is more common than overcounting (false positive); and (3) at most 1 nestling is missed/invented. The last rule was especially important to limit the number of cells in **p** that need to be estimated (reducing it from 16 to 10); it may be relaxed in theory but that will increase the necessary sample size to estimate all values in **p**; moreover, not fixing probabilities of highly unlikely combinations of *z* and *y* (e.g., observing 4 nestlings in a nest that has 1 nestling) may cause numerical problems. Based on this parameterization, we addressed three questions:

#### Question 1: How does the sample size of a paired data set affect estimates of p?

In paired data sets, we have direct measures of *z*, and we therefore expect unbiased estimates of **π** and *λ*, with precision increasing with sample size. We expect a much stronger impact of sample size on estimates of **p**, because sample size per element of **p** is much lower than for **π**. Also, how often certain nestling numbers are observed is affected by how common they are in the population, as well as the classification process; thus, for several cells in **p**, we may have 0 observations even at intermediate sample size. We therefore expect small sample bias in **p**, particularly in cells referring to rare states, that will decline with increasing *n*.

We generated 100 paired data sets for *n* = 10, 25, 50, 100 and 250 nests, mirroring a range of sample sizes in red kite monitoring programs (Sergio et al. 2019). We did so for each of the three alternative levels of fecundity. We estimated **π**, *λ* and **p** from the resulting data sets using our nestling count model and compared estimates to truth. We focus on absolute bias in estimates of **p** and relative bias and precision based on the coefficient of variation (CV = estimate divided by standard error) in estimates of *λ*. We report absolute bias for **p** and abstain from reporting a CV because several true values of **p** are fairly small, leading to extreme relative bias and CV values.

#### Question 2: How much precision do we gain in estimating fecundity when combining a small paired data set with uncertain counts, when both come from the same population (i.e., share fecundity)?

Climb counts are logistically and financially challenging to implement at large spatial scales; representative sampling of a population may require large sample sizes that are unfeasible with climb counts. It may therefore be beneficial to combine a fairly small set of paired counts with a larger data set consisting only of uncertain ground counts. In this scenario, again, because direct information on *z* is available from the paired data, we expect **π** and *λ* to be unbiased, but we expect precision to increase as we include additional uncertain counts.

We created combined data sets by combining paired data sets with *n* = 25 with uncertain count only data (generated from the same values of **π** as the paired data) of size *n*.*new* = 10, 25, 50, 100 or 250 (100 combined data sets for each value of *n*.*new*). We estimated **π**, *λ* and **p** from the resulting data sets and compared estimates to truth, as well as to estimates from the corresponding scenario from Question 1 (i.e., with *n* = 25 paired counts and no additional ground counts). We did so for all three levels of fecundity.

#### Question 3: How accurately can we estimate fecundity for a set of uncertain counts when we combine it with paired data sets of different sample sizes from a population with different fecundity?

This is likely the most important aspect of this model, as it would allow using paired counts from one population to estimate **π** and *λ* in another population with differing fecundity while assuming the classification process is the same.

We created combined data sets by combining paired counts (generated from baseline **π**) of size *n* = 10, 25, 50, 100 or 250 (i.e., 100 combined data sets for each value of *n*) with uncertain-only counts of size *n*.*new* = 50 generated from **π. new**. We did so separately for **π. new** = **π**_*high*_ and **π. new** = **π**_*low*_. We estimated **π. new**, *λ. new* and **p** from the resulting data sets and compared estimates to truth.

For all scenarios/questions, we further compared bias in model-estimated *λ* to bias if we calculated *λ* directly from uncertain counts (*λ*_*y*_), i.e., ignored any observation bias.

### Model implementation

We implemented the model in a Bayesian framework, for the ease with which this framework handles latent state variables. Specifically, we used the package nimble v.0.12.2 (de Valpine et al. 2017, 2022) in R v.4.2.1 (R Core Team 2022). To improve convergence of MCMC chains, we used the log-linear formulation of the multinomial model for both **π** and **p**, which estimates *log*(*γ*), then rescales to multinomial cell probabilities using 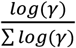. We wrote a custom Nimble function to fix some values for **p** to 0. We used moderately informative Normal priors (*μ* = 0, *σ* = 2) on *log*(*γ*) to ensure chain convergence. Note that when estimating only **π** from climb counts, convergence was not an issue and informative priors were not necessary, but extending the model to estimate **p**, even only from paired data, led to convergence issues in the absence of these moderately informative priors. For all models under Question 1 and 2, we ran 3 parallel chains with 5000 iterations per chain, discarding the first 2500 as burn-in; for Question 3, we ran chains for 10000 iterations, discarding the first 5000. We checked convergence by calculating the Gelman-Rubin statistic (Brooks and Gelman 1998) and treated chains as converged when 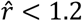 1.2 (this is a little less conservative than the typical cutoff of 1.1; for time constraints we wanted to avoid running a large number of models for longer, but for applications of the model to real data we would urge users to run the model long enough to obtain 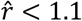). We also visually inspected trace plots of every 20th simulation iteration per scenario. We excluded non-converged simulation iterations from summary calculations and plots. The numbers of excluded simulation iterations are provided in Appendix 1: Tables S1-3.

### Problematic datasets

Traceplot inspections showed that for datasets in which some true state category was not represented at all in the paired counts (henceforth, datasets with missing truths), some values for **p** were not estimable (posteriors were flat over the 0-1 interval). This is intuitive: the model cannot estimate classification probabilities for a state not represented in the data. We therefore removed such data sets, leading to a lower number of simulation iterations: for *n* = 10, 70-75% of data sets had to be excluded; for *n* = 25, 20-45%; for *n* > 25, <10% of data sets had to be excluded. Because not all state categories are equally likely to be missing, however, this led to a biased sample of retained datasets, particularly for *n* = 10. For baseline *λ* and *λ*_*low*_, *z* = 4 was the rarest category and retained datasets (which had at least one nest with *z* = 4) therefore had an average number of nestlings higher than *λ* or *λ*_*low*_. For *λ*_*high*_, *z* = 1 was the rarest category and retained datasets (with at least one nest with *z* = 1) had a lower average number of nestlings than *λ*_*high*_. Only for *n* = 10 and 25 did this bias in retained data exceed 1% (Appendix 1: Table S4). To assess bias in model estimates of *λ* fairly for *n* = 10 and 25, we compared estimates against the retained sample mean 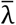 instead of the original true input value of *λ*.

Because dataset removal due to missing truths was applied only to paired counts, there is no systematic bias in additional uncertain count data, which were generated independently to address Question 2 and 3. For Question 2, however, the model assumes *λ* to be the same in the paired and the uncertain-only counts, which is not the case when datasets with missing truths are excluded from the paired data. To avoid this mismatch in *λ* between paired and uncertain count data in Question 2, we simulated these additional uncertain counts using the sample average values of 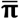 of retained datasets and compared estimates of *λ* from combined paired and uncertain counts against the retained sample mean 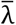. For simplicity, we still refer to scenarios in Question 1 and 2 by the original data-generating input values of *λ* (i.e., 2.0, 2.4 and 2.8).

### Population projections

To investigate how bias in fecundity affects population projections, we implemented a second simulation, projecting an age-structured population 50 years into the future, considering females only. Input parameter values for starting age distribution, age-specific proportion of the population that breeds, proportion of breeders that produce any fledglings, and age-specific survival were taken/adjusted from Sergio et al. (2021) and Mammen et al. (2017) and are summarized in Appendix 1: Table S5. We used two sets of age-specific survival probabilities, one that led to a growing population and one that led to a declining population. We combined each with values of **π** from the parameter estimation simulation to generate the total number of nestlings. Assuming a 50:50 sex ratio at nestling stage, we then randomly assigned each nestling a sex and only considered females going forward in time, to keep with the female-only projection. We projected the population with the true values of **π** (high and low), the corresponding expected observed values of **π** if we used ground counts only and ignored observation bias (**π**_*y*_), and the average (across simulation iterations) model-estimated values of 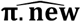 from Question 3, combining 50 uncertain counts generated from **π. new** with 50 paired counts generated from baseline **π**. Using the average over simulation iterations ignores the fact that estimates at each individual iteration were likely more different from truth than the average. But our interest here is to see the general impact on population trajectories when ignoring observation error vs when accounting for it; thus, we ignored variability in estimates in our projection. We used a starting population size of 100 females and we projected the population 1000 times over 50 years. We calculated average trajectories and 95% confidence intervals over the 1000 iterations, as well as average difference in population size in year 50 between the three projections using true, observed uncorrected, and estimated fecundity.

## Results

### Empirical data

The average number of young observed in climb counts at the 26 surveyed nests was 2.15 (range: 1 to 3); the average number based on ground counts for the same nests was 2.04 (range: 1 to 4), thus showing approximately 5% negative bias. At 19 nests, ground counts yielded the same values as climb counts; climb counts were biased low by one nestling in 5 and biased high by one nestling in 2 nests. All combinations of ground and climb counts are available along with the code to implement simulations.

### Simulation - parameter estimation

Input parameter settings of nest probabilities (**π**) and classification probabilities (**p**) led to 7% negative bias in fecundity when ignoring observation bias for λ and *λ*_*high*_, and 6% negative bias for *λ*_*low*_.

For Questions 1-3, median bias across all kept simulation iterations, CV of *λ*, as well the RMSE of all parameters for all scenarios can be found in Appendix 2.

#### Question 1

As expected, when analyzing paired counts only, estimates of *λ* were essentially unbiased, with relative bias < 2% for all scenarios (compared to retained sample mean for *n* = 10 and 25, compared to true input value for all other *n*; Appendix1: Figure S1A) and the CV of *λ* declined with increasing sample size, but was already low at *n* = 10 (ranging from 8.9% for *λ*_*high*_ to 11.4% for *λ*_*low*_; Appendix1: Figure S1B). Estimates of **π** showed mild bias at *n* = 10 and 25, but were essentially unbiased at larger sample sizes (Appendix1: Figure S2). As expected, estimates of **p** showed considerable bias for small sample sizes (*n* = 10 and 25) and were only largely unbiased at *n* = 250, with some bias remaining in some of the probabilities (Figure 1 for *λ*, Appendix1: Figure S3 for *λ*_*low*_ and *λ*_*high*_).

**Figure 1.**
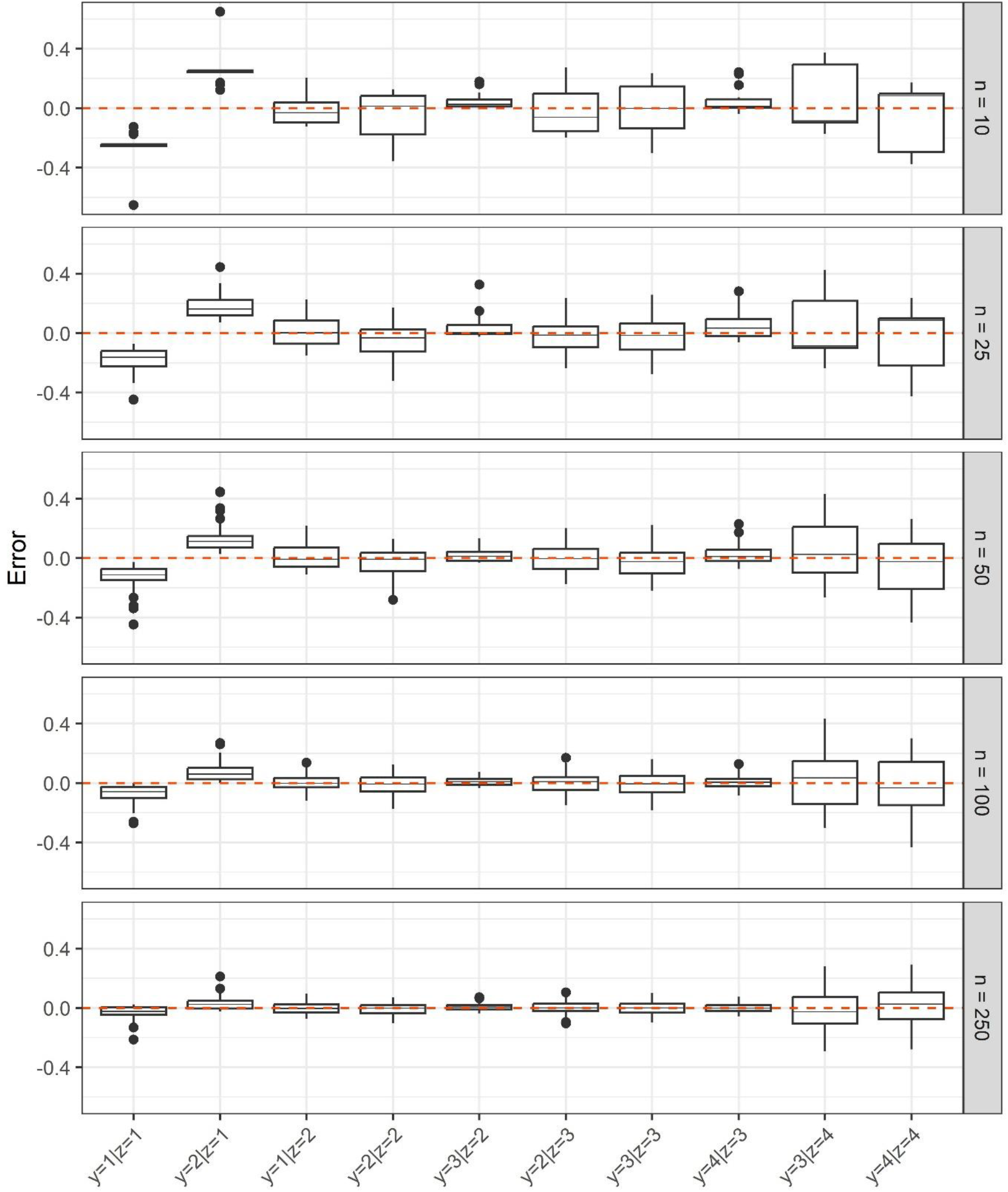
Absolute error (boxplots) and bias (median line) of estimates of classification probabilities **p** of observing *y* young when the true number of young is *z* from paired certain and uncertain counts of different sample sizes (*n*) from a model accounting for over- and undercounting of young. Red dashed line indicates 0 error/bias. Values of **p** for missing combinations of *y* and *z* were fixed to 0.

#### Question 2

Adding uncertain ground counts *y*.*new* to 25 paired counts, with both datasets having the same underlying fecundity, bias in *λ* was reduced with 10 additional ground counts; adding more ground counts slightly increased bias, though it remained very low overall (< 1.6%; Figure 2A). Bias in *λ* was generally negative. Adding ground counts marginally improved the CV of *λ*, e.g. by 2-3% comparing 250 to 0 additional ground counts (Figure 2B). Overall, patterns of bias in **π** were similar to those in *λ*, with 10 additional ground counts improving on estimates, but more ground counts increasing bias again slightly (Appendix1: Figure S4). Similarly, bias in **p** (e.g., Appendix1: Figure S5) remained very similar to that observed with no additional ground counts, as expected, since *y*.*new* does not contribute information to estimating **p**. Interestingly, the more ground counts were added, the higher the rate of non-convergence (Appendix 1: Table S2); this was not the case in Question 3.

**Figure 2.**
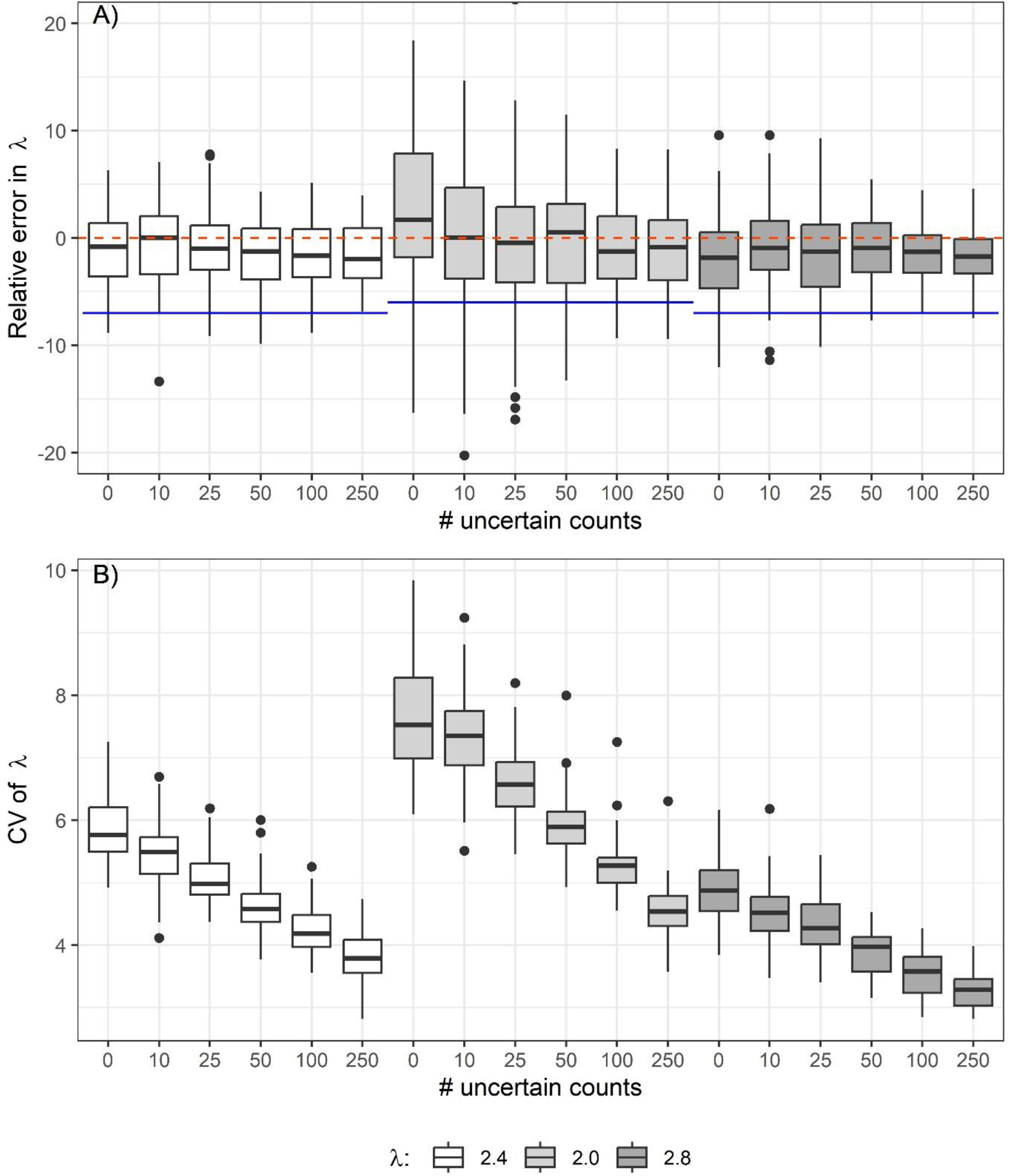
Relative error (boxplots) and bias (median) (A) and coefficient of variation (CV; B) of estimates of fecundity *λ* (average number of young per nest) from a model accounting for over- and undercounting of young, fit to 25 paired certain and uncertain counts plus varying numbers of additional uncertain ground counts (x axis), when both data sets come from the same population (with the same fecundity; Question 2). Red dashed line and blue lines in A show 0 bias/error and relative bias if fecundity was calculated based on uncertain-only counts, ignoring observation bias, respectively.

#### Question 3

When combining paired counts of different sample sizes *n* from a population with nest probabilities **π** with 50 uncertain counts from a population with **π. new**, estimates of *λ. new* were largely unbiased across all sample sizes (Figure 3). There was a tendency for bias to increase with increasing *n* for *λ. new* = *λ*_*low*_, however, even at *n* = 250, bias was only 2%. Whereas in Question 1 all values of **π** became essentially unbiased at *n* = 50 or 100, here, some bias in **π. new** remained at all sample sizes, with no obvious trends (Appendix1: Figure S6). Bias in estimates of **p** decreased with increasing paired sample size; estimates of **p** also became increasingly similar to those under Question 1 with increasing sample size (Appendix 1: Figure S7).

**Figure 3.**
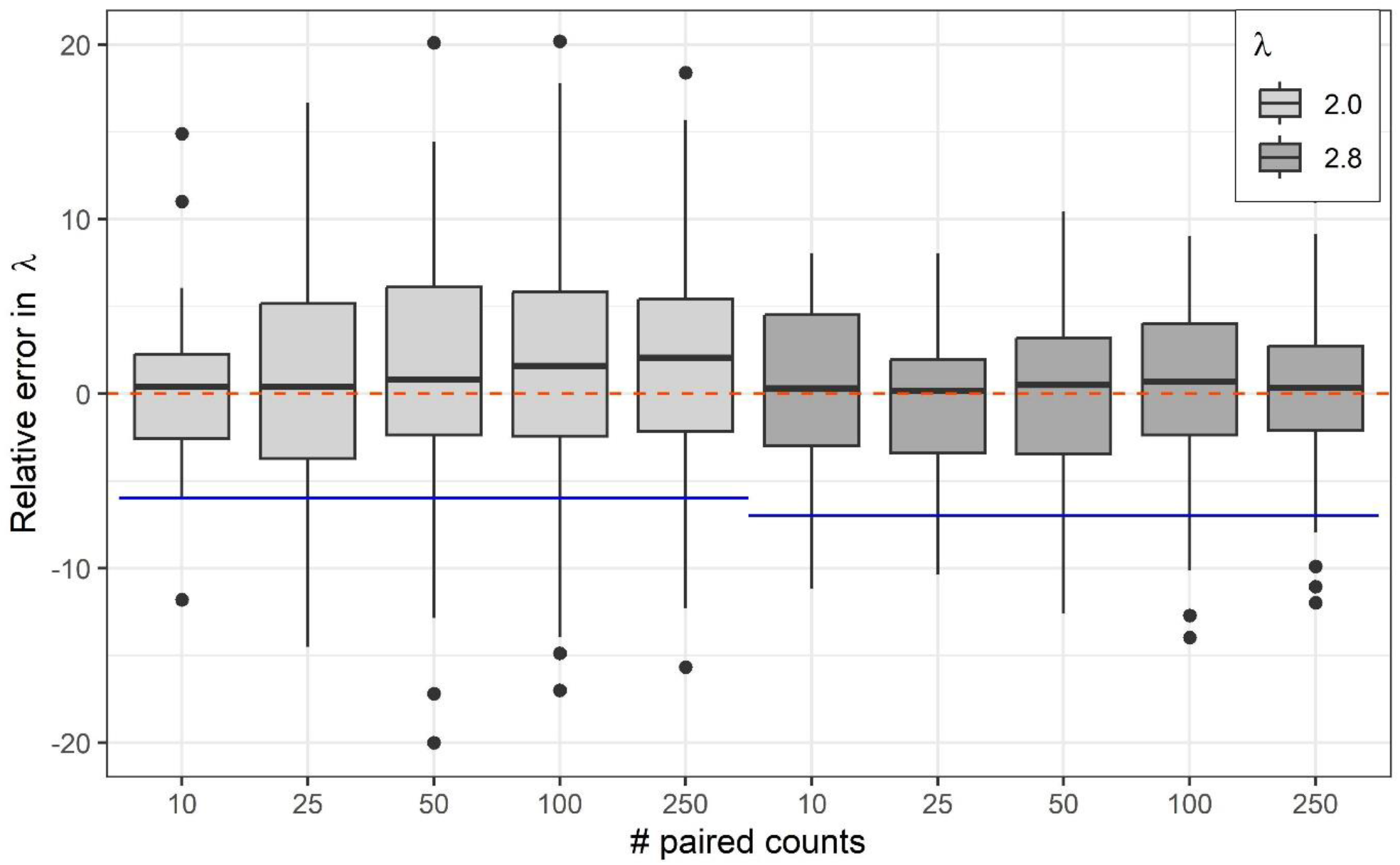
Relative error (boxplots) and bias (median) of estimates of fecundity *λ* (average number of young per nest) from a model accounting for over- and undercounting of young, fit to 50 uncertain counts from a population with *λ* = 2 or *λ* = 2.8 plus varying numbers of paired certain and uncertain counts (x axis) from a populations with *λ* = 2.4. Red dashed line shows 0 bias/error, and blue lines show bias if fecundity was calculated based on uncertain counts only, ignoring observation bias.

### Population projections

Using nest probabilities based on uncertain-only (i.e., ground) counts, ignoring observation bias, led to negative bias in projected population trajectories. That bias was stronger in growing populations and at higher fecundity (**π**_*high*_), because of the exponential growth underlying the projections. In agreement with the largely unbiased estimates of fecundity from the parameter estimation simulations, using average model-estimated nest probabilities led to population trajectories that were largely identical to those using true input values (Figure 4). Consequently, relative bias in population size in year 50 was negligible when using estimated nest probabilities (< 1% to 5% negative bias), but considerable when not accounting for observation bias (25% to 42% negative bias; Table 1).

**Table 1.** Percent bias in projected population size at time *T*=50 when using the observed uncertain counts of nestlings to determine fecundity (observed) and when using estimated fecundity accounting for observation error in uncertain counts (estimated). Median (2.5^th^ and 97.5^th^ percentile) across 1000 simulated population trajectories, relative to the trajectory using true fecundity for increasing and decreasing populations with either high (input π_*high*_) or low (input π_*low*_) fecundity.

| Population | Observed vs true | Estimated vs true |
| --- | --- | --- |
| High, increasing | -38.91 (-63.52 - -10.87) | -3.03 (-41.51 - 42.11) |
| High, decreasing | -32.69 (-87.06 - 50.21) | -9.4 (-76.7 - 104.5) |
| Low, increasing | -28.92 (-66.09 - 22.23) | 4.73 (-49.94 - 77.16) |
| Low, decreasing | -31.7 (-100 - 127.66) | -8.94 (-100 - 195.96) |

**Figure 4.**
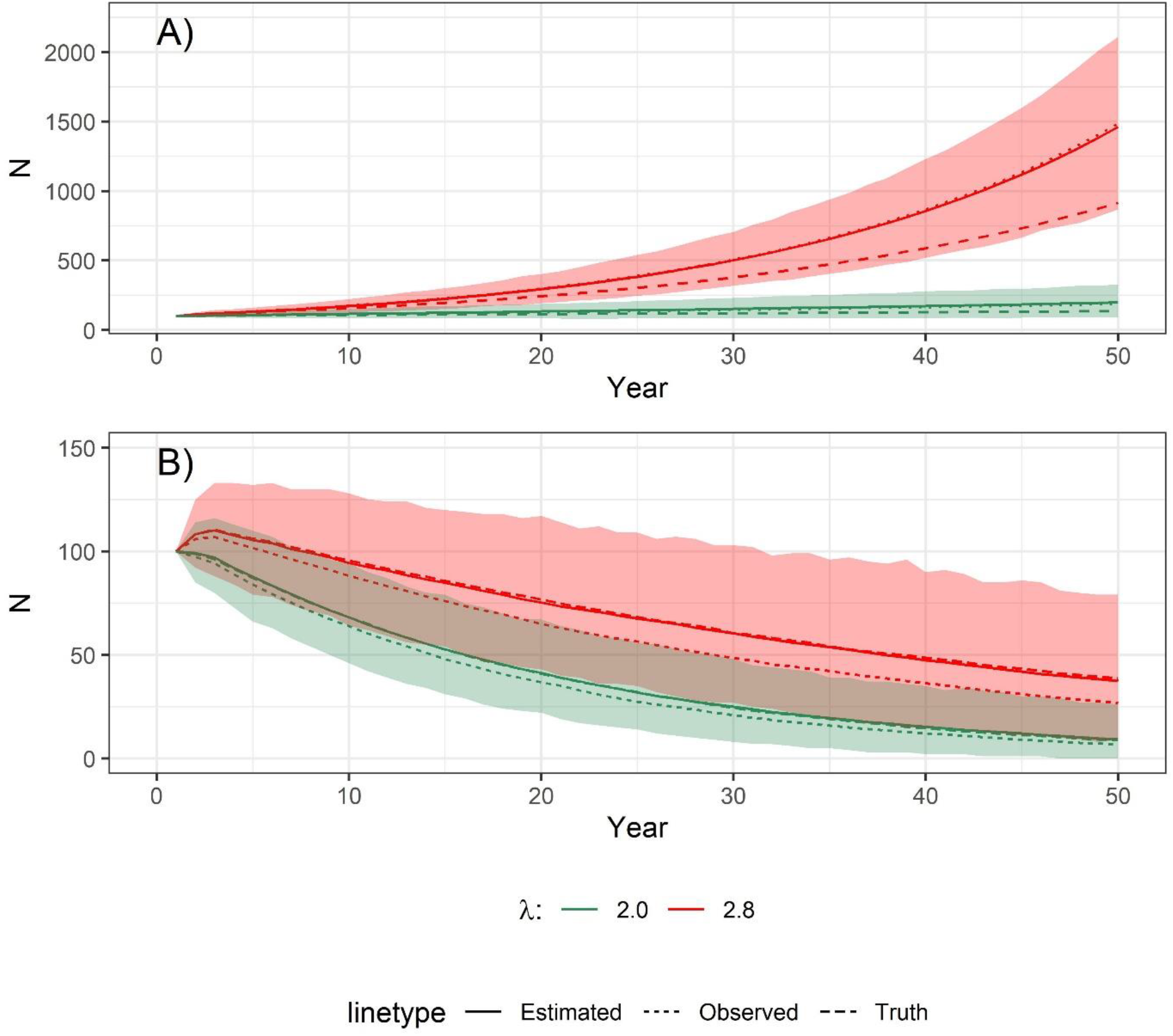
Population projections for growing (A) and declining (B) populations with two levels of fecundity (*λ*). Solid lines show median trajectory and colored polygons 2.5^th^ and 97.5^th^ percentiles of 1000 projections using true input value of *λ*. Dashed lines show median trajectory (across 1000 projections) when calculating *λ* based solely on uncertain counts of young, and dotted line shows median trajectory (across 1000 projections) when estimating *λ* with a model accounting for over- and undercounting of young.

## Discussion

During a count-based survey, inadvertently counting the same individual more than once (i.e., a form of false positive error) is a common problem (e.g., in camera trapping - McKibben et al. 2023; unmanned aerial surveys - Brack et al. 2018). Yet, few modeling approaches explicitly account for such false positives (but see Kéry and Royle 2015). In our classification model, we treat the true state - the number of nestlings - as categorical. This allowed us to adopt the strategies developed for multi-state occupancy models that account for false positives and state uncertainty (Royle and Link 2005, 2006, Miller et al. 2011). In particular, we used a calibration approach that combines uncertain with certain counts to learn about the probabilities of observing a certain number of nestlings conditional on the true number of nestlings, **p**, and then using those probabilities to correct uncertain counts in a separate dataset (Chambert et al. 2015).

We found that our approach produced largely unbiased estimates of fecundity for nestling counts suffering from both false positive and false negative observation error. Unbiased estimates persisted regardless of whether the separate dataset had the same data-generating value of fecundity (i.e., came from the same biological population) or a different value (i.e., came from a different population). Ignoring observation errors in the number of nestlings led to underestimating forecasted abundances, particularly in large or growing populations. Using estimates that corrected for both false negatives and false positives, however, led to population trajectories that closely resembled true trajectories.

The most crucial variable determining the performance of our classification model was the sample size of the paired counts (i.e., the number of nests for which both certain and uncertain observations were available). Specifically, some classification probabilities **p** were not estimable when a dataset of true nestlings was missing a nestling category (e.g., no nest in the dataset had 4 nestlings), as in those cases the dataset contains no information on the classification probabilities related to the missing true state. Missing true states were, of course, more common in small data sets; in our simulation, a paired *n* = 50 was generally sufficient to avoid this problem. The necessary sample size, however, depends on the distribution of probabilities, **π**, among the true states. In particular, the rarer one or more categories become, the larger the sample has to be to detect all categories.

A truncated Poisson distribution is a potential alternative for modeling nest probabilities, **π**. With a single parameter (as opposed to *Z*-1 parameters in the multinomial model), it is more parsimonious and its specific properties can inform nest probabilities for categories with no observations. We found, however, that a truncated Poisson did not fit the empirical data well and led to biased estimates of **π**. Limited trials suggested that this poor fit increased bias in estimates of fecundity when combining paired data with uncertain counts from populations with differing fecundities. A more thorough evaluation is necessary to determine the extent of this bias as a function of the degree of lack of fit.

As an alternative, unobserved or very rare state categories could be excluded when estimating fecundity; that has the added advantage of reducing the size of the classification matrix **p**. We did not explore any potential effect of excluding non-represented nestling categories on bias and precision in estimates of fecundity. Even in datasets without missing true categories, many classification probabilities showed large bias; in individual simulation iterations, some probabilities were often barely estimable, with large uncertainty, and we had to use mildly informative priors on the log-probabilities of the classification model to ensure chain convergence. All of these issues are likely due to small sample sizes for individual classification probabilities, even though we already reduced their number by limiting the difference between true states and observations to 1. These limitations highlight that with an increasing number of true states, the size of the classification matrix becomes unwieldy and parameters comprising this matrix become challenging to estimate (Royle and Link 2006). Other constraints, such as imposing that true classification probabilities are higher than false classification probabilities (Royle and Link 2005, 2006, Miller et al. 2011), and that undercounting is more likely than overcounting are justified based on our empirical data and could help further improve the performance of the model we developed. Still, based on our simulation results we believe this classification model is of limited use for species with a wider possible range of brood sizes than what we considered here.

There are ways in which the classification model could be simplified. For example, the imperfect observations could be regarded as Poisson random variables with mean *z* × *ϱ*, where *ϱ* is a detection rate correcting for net bias in nestling counts (for example, in our empirical data, undercounts were more frequent than overcounts, thus, net bias was negative and *ϱ* would be < 1). This is akin to the observation process in the Poisson-Poisson N-mixture model (Kéry and Royle 2015), which allows for both over and undercounting. Preliminary trials with our simulated data suggest that this model will lead to bias in estimates of fecundity when paired counts are combined with uncertain-only counts, even though the paired data provide direct information on *ϱ*. N-mixture models rely on temporally replicated sampling, whereas our data has a single observation per nest. When it is easier to obtain repeated uncertain counts from nests than to climb them to ascertain the true number of nestlings, a PP N-mixture model may be a viable alternative to our classification model. The standard PP N-mixture model has been shown to provide largely unbiased estimates of abundance (Nakashima 2020). How well a modified version with a multinomial (or truncated Poisson) state process would perform for estimating fecundity remains to be evaluated.

Our empirical data suggested that both over and undercounting misses the true number of nestlings by at most 1. If that is the case, the classification process can be broken down into two Bernoulli processes: whether the number of nestlings is counted correctly (with probability *p*_*c*_), and if not, then whether the number is overcounted (with *p*_*over*_; as opposed to undercounted). The classification probabilities can then be written as a function of these Bernoulli probabilities. If either or both probabilities can be assumed to be the same regardless of the true state, this will reduce the number of observation parameters that need to be estimated. Initial trials, using our original simulated data and allowing *p*_*c*_ to vary by true state but fixing *p*_*over*_ suggested that this simplified model may yield similarly unbiased estimates of fecundity as the more complex data generating model (though in our case, the two model versions differed only by one free parameter).

Adding uncertain counts from the same biological population (i.e., with the same fecundity) to 25 paired counts (Question 2) only marginally improved the precision in estimates of *λ* and led to more convergence problems. Thus, in a study focusing on a single population with constant fecundity, there is little to no benefit from investing effort in additional ground counts. All effort in such a case should be dedicated to collecting unbiased brood size information at as many locations as possible. In contrast, adding even a small number of paired counts to a dataset of uncertain counts from a population with different fecundity greatly reduced bias in estimates of *λ*_*new*_ compared to ignoring observation bias altogether. As long as the observation process (i.e., classification probabilities) can be assumed to be the same in different populations, information on these probabilities from one population can be leveraged to obtain unbiased estimates of reproduction in another population.

The validity of detection parameters beyond the spatial and temporal scope of sampling used to estimate them needs to be evaluated carefully (e.g., Williams et al. 2002). An important future step is therefore to identify factors affecting classification probabilities and to integrate them into the classification model. For example, nest visibility (and hence, the probability of correctly identifying the number of nestlings) may be higher in deciduous than in mixed or coniferous forests. Identifying important predictors of classification probabilities would make estimates of **p** more transferable to other populations, because sources of variation in the classification parameters could then be taken into account. Attempts to do so with the present dataset failed because of the small sample size.

Alternatively to count-based methods to estimate brood size or fecundity, capture-mark-recapture methods can be used to estimate age-specific population sizes and derive fecundity as number of juveniles per adult (e.g., Sen and Akçakaya 2022). Capture and recapture of individuals, however, requires significant expenditures of time and money by investigators (Lieury et al. 2017). Such repeated sampling may thus not be feasible in many situations.

### Recommendations for data collection

There are several important considerations when obtaining data necessary for applying our modeling framework, first and foremost that it requires at least some paired data that includes certain and uncertain counts. In situations where nests are unaccessible to humans, drones may prove to be a suitable approach to obtain certain counts. In fact, they may generally be a cheaper and safer alternative to climbing nest trees if counting nestlings is the sole purpose, allowing for larger paired samples. However, accuracy of drones in counting nestlings has not been assessed yet.

Unless researchers adapt this framework to incorporate covariates for the classification probabilities, potential biases should be minimized by keeping the observation conditions as consistent as possible. These include weather, time of day, nestling age, duration of observation, method of observation (e.g., binocular, spotting scope), and distance from the ground observer’s position to the young. Fuller et al. (1995) recommended observing the nest for 30 to 60 minutes from the ground to accurately count the young. Observing young for > 1 hour likely increases disturbance and may increase the risk of overcounting the number of nestlings.

The paired ground and climb counts should be implemented during a single visit when at least one observer is present at all times, to ensure no young are lost (e.g., due to predation) between the two observations. A possible exception is when ground counts are paired with perfect counts obtained via video documentation of the nest. Installing a camera would require climbing to the nest at an earlier stage, disturbing both parents and nestlings when they are more vulnerable (Götmark 1992, Ibáñez-Alamo et al. 2012). Cameras could be installed during the non-breeding season at previously used nests but without guarantee that these will be occupied again.

### Limitations of this study

The present study has several limitations. First, the observed bias in uncertain ground counts in the example data set – and therefore, in the simulated data – was very low (6-7%) and model estimates of fecundity would likely always contain biased estimates based on uncertain counts within their confidence intervals (i.e., statistically, model estimates and biased estimtes would be deemed not statistically different from each other). Especially in long-lived species, whose population trajectories are less sensitive to recruitment (Sæther et al. 2013), the bias considered here may not be ecologically relevant. Our model is therefore likely more useful in situations where observation bias is stronger than what we considered here, or when studying short-lived species.

In addition, our model evaluation of bias and precision was based on a setting in which all nests had at least one nestling present and observed. By exluding the 0 state and respective observation categories, we reduced the number of parameters to be estimated by the model. In many real world applications, it will be necessary to include the 0 categories, thus risking even more estimability and identifiability issues than we encountered in our simulation. We illustrate through model evaluation how to address these difficulties by making restrictive assumptions about state–observation combinations (i.e., fixing certain classification probabilities to 0) and discuss other constraints (e.g., Royle and Link 2005, 2006, Miller et al. 2011) that could be employed in such situations.

Finally, the model is not only data-hungry but may also be dificult to implement for some practitioners. Alternatively, obtaining unbiased estimates of fecundity can also be achieved by foregoing a formal model altgether and simply calculating a correction factor *c* from all paired counts as 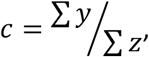 then use *c* to adjust observations from uncertain-only counts to obtain an unbiased estimate of *λ* (Appendix 1: Figure S8). This approach, however, does not allow for the inclusion of covariates affecting classification probabilities (or here, the correction factor), which we believe is an important extension to make our model more applicable under real field conditions.

### Conclusion

Fecundity is a key parameter affecting population dynamics (Sibly and Hone 2002), and we showed that even apparently low bias in fecundity estimates can lead to considerable bias in population projections under certain conditions. This can have implications for species conservation, for example, when conducting a population viability analysis (Boyce 1992). This highlights the need for statistical models that can produce unbiased estimates of fecundity taking into account both false positive and false negative observation bias. We believe the model presented here represents a useful approach to dealing with both kinds of observation bias while taking into account the specific characteristics of nestling count data, particularly, the limited number of true and observed states. The approach hinges on the assumption that the classification process is the same for data with and without certain observations. To improve transferability of classification probabilities between points in space or time, we believe the most important next step is to examine potential predictors of these probabilities (e.g., time spent observing), and to extend the model to incorporate them. Including such covariates would be easier if the number of classification parameters could be reduced and we sketch out several possibilities for such simplified models, the behavior of which under different scenarios remains to be explored. More broadly, our approach contributes to the field of models accounting for biases induced by false-positive observations, which remains underrepresented in the literature even though its importance has been highlighted repeatedly.

## Appendices

## Appendix 1

Additional information and results from estimation simulation and population projections

## Appendix 2

Median bias, precision and accuracy of parameter estimates

Appendices can be found at doi.org/10.5281/zenodo.10245397 (Sollmann et al. 2024).

## Acknowledgements

We thank R. Raab for allowing the student assistants to join the telemetry team in visiting the nests for data collection. F. Knufinke, M. Kolbe, E. Schöll and two reviewers provided useful input in the development of the manuscript. We thank M. Malvezin for assistance with refining the field sampling protocol and data collection.

## Data, scripts, code, and supplementary information availability

Data, scripts and code can be found at doi.org/10.5281/zenodo.10245802 (Sollmann 2024).

